# *Microcystis aeruginosa* grown in different defined media leads to different cultivable heterotrophic bacteria composition that could influence cyanobacterial morphological characteristics and growth properties

**DOI:** 10.1101/721175

**Authors:** Nicholas M.H. Khong, Yam Sim Khaw, Muhammad Farhan Nazarudin, Fatimah Md. Yusoff

**Affiliations:** Institute of Bioscience, Universiti Putra Malaysia, UPM Serdang, 43400, Selangor Darul Ehsan, Malaysia; Department of Aquaculture, Faculty of Agriculture, Universiti Putra Malaysia, UPM Serdang, 43400, Selangor Darul Ehsan, Malaysia; International Institute of Aquaculture and Aquatic Sciences, Universiti Putra Malaysia, UPM Serdang, 43400, Selangor Darul Ehsan, Malaysia

**Keywords:** Cyanobacteria, *Microcystis*, non-microcystin production, heterotrophic bacteria, morphology, growth, bacterial-microalgae interaction, axenic culture

## Abstract

Cyanobacterial blooms involving *Microcystis* spp. often pose severe problems to the environment and general community due to their persistent presence in eutrophic water bodies and potential to form blooms. Bacterial associations are known to alter microenvironment of *Microcystis* and potentially influence their development. This study aimed to study cultivable heterotrophic bacteria composition that developed symbiotically with *Microcystis aeruginosa* naturally as well as those cultured under defined media and their possible effects on the morphology and growth properties of the cyanobacterium. *M. aeruginosa* (UPMC-A0051) was isolated during a bloom from Putrajaya Lake, Malaysia and characterized as a non microcystin-producing cyanobacterium using PCR and chromatographic methods. Associated heterotrophic bacteria were then isolated and identified from the culture media as well as the lake where the cyanobacterium was originally isolated. A total of 16 bacterial species were isolated from the lake and none of them were similar to the bacteria associated with *M. aeruginosa* cultured in artificial media. Cultivable heterotrophic bacteria composition associated with *M. aeruginosa* were also distinct in different culture media, despite the same inoculum. These bacteria were classified under *Actinobacteria, α.-Proteobacteria* and *β-Proteobacteria*. Under different bacterial associations, *M. aeruginosa* cultivated in defined media showed different colony morphology and growth properties. The present study demonstrated that distinct bacterial composition observed in different culture media could be responsible for dissimilar cyanobacterium morphology and growth rate, particularly on the clustering pattern. In the axenic culture, the growth of *M. aeruginosa* was significantly reduced indicating the influence of associated bacteria on the development of cyanobacterial colonies.

## Introduction

Episodes of cyanobacterial blooms have been reported unceasingly worldwide over the past century due to increased unsustainable anthropogenic activities resulting in related incidences of water enrichment and eutrophication. The blooms affect fishery resources, recreational activities, domestic water usage and aesthetic values of aquatic ecosystems. Previous studies of cyanobacterial water blooms have focused on cyanobacterial species and various factors in influencing the cyanobacterial growth [1].

*Microcystis* frequently forms dense surface blooms in freshwater environments leading to serious water quality problems. It has been established that cyanobacterial blooms in temperate waters are significantly different from those in tropical waters [2]. At such, algae reinvasion is a major interest in the study of temperate *Microcystis* blooms which have been reported to be influenced by environmental factors. The dominance of *Microcystis*, like many other species of cyanobacteria is mainly due to their wide range of morphological and physiological adaptive mechanisms especially their buoyancy properties, nitrogen-fixing capacity, colony formation and aggregation characteristics [3], which provide them with competitive advantages over other microalgae species. However, with practically no major seasonal fluctuations in the tropics, the occurrence, persistence and dominance of *Microcystis* blooms and their toxin production in tropical waters remain relatively understudied. It may be possible that *Microcystis* blooms and their toxin release are triggered by microbial interactions within the water body. It has been shown that associated bacteria are responsible for the induction of aggregation and colony formation of *Microcystis* [4].

Some studies have demonstrated significant effects of bacteria-microalgae interactions on microalgae growth and even morphological changes [4-6]. In natural environment, interactions between *Microcystis* and its associated bacteria could be classified as mutualism, commensalism and exploitation. One example of mutualism involves the inter-support of heterotrophic bacterial growth and *Microcystis* via production of complimentary byproducts i.e. organic matter synthesized by *Microcystis* and microalgal growth factors (e.g. vitamin) by bacteria [7]. Commensal bacteria degrade microcystin from *Microcystis* cells possibly for its essential nutrients, whereas *Microcystis* cells are not affected [8]. On the other hand, algicidal effect of bacteria on *Microcystis* cells is an example of exploitation [9].

*Microcystis* species are increasingly cultured in controlled environments for the study on applications of their secondary products, byproducts and toxins. As a general observation, laboratory cultured microalgae, including *Microcystis*, will differ appreciably from when they are obtained from the environment, where many would attribute to the difference in biochemical parameters between the culture media and the algae’s natural habitat. It is hypothesized that different culture media composition would lead to unique cultivable heterotrophic bacteria composition that might be the key factor in affecting the colony morphology and growth properties of *Microcystis*. Currently, the link between *Microcystis*-associated bacteria in different defined culture media with *Microcystis* still remain unclear. The purpose of the study is to evaluate a tropical *Microcystis* sp. and its associated cultivable heterotrophic bacteria composition from two most commonly used microalgal culture media and the possible role of these bacteria in shaping the morphology and growth properties of *Microcystis* sp.

## Materials and methods

### Algal collection, isolation and culture

Water samples were collected from Putrajaya Lake, Malaysia (2.9419° N, 101.6891° E) from July 2015 to October 2015 using 35 μm mesh size plankton net. The samples were transported to the laboratory in sample bottles with loose cover and used for isolation of *Microcystis* sp. Two growth media i.e. BG11 medium (Concentration, M in final medium: NaNO_3_, 1.76 × 10^-2^; K_2_HPO_4_, 2.24 × 10^-4^; MgSO_4_.7H_2_O, 3.04 × 10^-4^; CaCl_2_.2H_2_O, 2.45 × 10^-4^; Na_2_CO_3_, 1.89 × 10^-4^; Na_2_EDTA.2H_2_O, 2.79 × 10^-6^; Ferric ammonium citrate, 3 × 10^-5^; Citric acid, 3.12 × 10^-5^; H_3_BO_3_, 4.63 × 10^-5^; MnCl_2_.4H_2_O, 9.15 × 10^-6^; ZnSO_4_.7H_2_O, 7.65 × 10^-7^; CuSO_4_.5H_2_O, 3.16 × 10^-7^; Na_2_MoO_4_.2H_2_O, 1.61 × 10^-6^; Co(NO_3_) _2_.6H_2_O, 1.70 × 10^-7^; pH 7) and Bold’s basal medium (BBM) (Concentration, M in final medium: NaNO_3_, 2.94 × 10^-3^; K_2_HPO_4_, 4.31 × 10^-4^; KH_2_PO_4_, 1.29 × 10^-3^; MgSO_4_.7H_2_O, 3.04 × 10^-4^; CaCl_2_.2H_2_O, 1.70 × 10^-4^; NaCl, 4.28 × 10^-4^; Na_2_EDTA.2H_2_O, 1.71 × 10^-4^; KOH, 5.53 × 10^-4^; FeSO_4_.7H_2_O, 1.79 × 10^-5^; H_3_BO_3_, 1.85 × 10^-4^; MnCl_2_.4H_2_O, 7.28 × 10^-6^; ZnSO_4_.7H_2_O, 3.07 × 10^-5^; CuSO_4_.5H_2_O, 6.29 × 10^-6^; Na_2_MoO_4_.2H_2_O, 4.93 × 10^-6^; Co(NO_3_) _2_.6H_2_O, 1.68 × 10^-6^; pH 7) were prepared aseptically. Isolation of *Microcystis* sp. was carried out by selectively picking the cyanobacterium using a modified micropipette followed by successive washings with sterile media under an Axioskop 2 light microscope (Carl Zeiss, Oberkochen, Germany). The washed cyanobacterium was streaked on BG11 agar plates for two times. The purity of the monoclonal cyanobacterium was checked regularly using light microscope. Once a single colony of the cyanobacterium was obtained, it was cultivated in BG11 and BBM media at 25°C ± 1°C under illumination of cool fluorescent light (2000 lux) with 12: 12 hour light and dark cycle.

### Morphology

The morphology of the isolated *Microcystis* sp. from the lake water and the cultivated *Microcystis* sp. was observed using a light microscope, attached with a charged couple device camera (Carl Zeiss, Oberkochen, Germany) under 100×, 200×, 400×, 1000× magnifications. *Microcystis* sp. was morphologically classified based on the characteristics described by Desikachary [10]. The diameter of each *Microcystis* cell (an average of 50 cells) was measured using a calibrated micrometer scale for three times to obtain the mean size.

### Molecular identification

Microalgal species was identified using 16S rRNA PCR. Prior to amplification, 1 mL *Microcystis* sp. culture (OD_680_: 0.1) was centrifuged for 10 min at 8000×g for DNA extraction. DNA extraction of *Microcystis* sp. culture was performed using FavorPrep Tissue Genomic DNA Extraction Mini Kit (Favorgen, Ping-Tung, Taiwan) following the manufacturer’s instructions. The 16S rRNA gene of the cyanobacteria strain was amplified using two sets of oligonucleotide primers, (1) CYA106F and CYA781R [11] and (2) 16S PSF and 16S R (universal bacterial primer, 1492R) [12]. In addition, pcbβF and pcαR primer sets were utilized to amplify the phycocyanin operon of the cyanobacteria [13]. PCR amplification was performed in a total 25 µL volume comprised of 1× Colorless GoTaq Flexi Buffer, 3 mM MgCl_2_, 0.2 mM dNTP (Fermentas, Waltham Massachusetts, USA), 1.0 μM of each forward and reverse primer, 2.5 U GoTaq Flexi DNA Polymerase (Promega, Madison USA) and 3 μL template. Thermal cycling profile was performed using an initial denaturation at 95°C for 5 min, followed by 35 cycles, each consisting of denaturation at 95°C for 30 s, annealing at 50°C (for 16S rDNA primer sets) or 55°C (for pcbβF and pcαR primer set) at 30 s and extension at 72°C for 1 min, and final extension at 72°C for 7 min using T-Personal Thermal Cycler (Biometra, Jena, Germany). Prior to sequencing, the PCR products were gel-purified using gel/ PCR DNA fragments extraction kit (Geneaid, New Taipei City, Taiwan). Sequencing of these purified products was performed in bi-direction with both forward and reverse primers by Apical Scientific Laboratories Sdn Bhd (Selangor, Malaysia) using ABI 3770 sequencer (Applied Biosystems, Foster City, USA).

### Microcystin-producing properties

#### PCR detection of microcystin gene

The microcystin primers, tox4f (forward primer) and tox4r (reverse primer), adopted from Kurmayer et al. [14] that target the *mcy*B A1 domain were used in this study. The previously extracted *Microcystis* sp. DNA was utilized as the PCR template. A total volume of 25 µL PCR reaction was prepared as above. The PCR thermal cycling protocols were an initial denaturation at 94°C for 10 min, followed by 35 cycles at 94°C for 30 s (denaturation), at 50°C for 30 s (annealing), 72°C for 1 min (extension) and final extension at 72°C for 7 min with the use of T-Personal Thermal Cycler (Biometra, Jena, Germany). As for direct sequencing, the amplified *mcy*B products were first gel-purified using gel/PCR DNA fragments extraction kit (Geneaid, New Taipei, Taiwan) as per manufacturer’s instructions followed by bi-direction sequencing involving both forward and reverse primers (First Base Laboratories Sdn Bhd, Selangor, Malaysia).

#### Liquid Chromatography Mass Spectrometry/Mass Spectrometry (LC MS/MS)

Dense growing *Microcystis* cultures from two media were centrifuged at 5000×g for 15 min and pellets were lyophilized and stored at −20°C prior to analysis. Extractions of microcystins were performed by sonicating 0.1 g lyophilized *Microcystis* cells in 70% methanol (w/v) for 20 minutes. Three similar extractions were conducted and pooled together and then, the residue was evaporated at 60°C. Methanol solution (10%) was used to redissolve the residue. The extract was fractionated using a preconditioned ODS solid phase extraction (SPE) cartridge (Merck, New Jersey, USA).

A step gradient of 30% to 100% methanol in water (v/v) was used to elute the extract. The 60% methanol (v/v) fraction was dried *in vacuo* and resuspended in 1 mL 100% methanol. Qualitative analysis of microcystin in the fraction was carried out by LC MS/MS analysis using analytical column, Acclaim PepMap100 C18 (150 × 1.0 mm, 3 µm; Thermo Scientific, Waltham, USA) with the mobile phase of water containing 0.1% formic acid and acetonitrile containing 0.1% formic acid [15]. The LC MS/MS system was an Ultimate 3000 x2 Dual Rapid Separation LC system (Dionex, Sunnyvale, USA) equipped with an electrospray ionization (ESI) interface. The gradient condition was set according to Zhang et al. [15] and the flow rate was set at 150 μL/min. Microcystin standard comprises of MC-RR, MC-YR and MC-LR, which eluted at 5.62, 6.85 and 6.93 minutes, respectively. The MS/MS spectrum of MC-RR is characterized by major fragment ions at *m/z* 452, 505 and 887, while MC-YR contains major fragment ions at *m/z* 599, 916, 1017. The MS/MS spectrum of MC-LR comprises of major fragment ions at *m/z* 553, 599, 866, 967.

#### Growth measurement

Unialgal *Microcystis* sp. culture was prepared by picking a single colony of *Microcystis* sp. from BG11 agar plate. To subculture from the original *Microcystis* sp. culture, the cells in exponential phase were utilized as inoculum. Growth measurement was made using cultures produced from the inoculation of 300 mL of the stock culture (BG11 and BBM specific) making up to 3 L with BG11 and BBM media yielding a final OD_680_ of approximately 0.1. All treatments were conducted in triplicates. The culture was gently agitated to ensure the culture uniformity before each growth measurement. Axenic cultures of *Microcystis* sp. were prepared using the procedures described previously [16-17]. Axenic condition was examined using epifluorescence microscope techniques [16-17].

*Microcystis* sp. growth assessment was performed by measuring biomass in dry weight as well as observation of optical density at 680 nm using a spectrophotometer (Shimadzu, Kyoto, Japan). Growth measurement for *Microcystis* sp. was conducted daily for 30 days. Specific growth rate (*µ*) was calculated according to the equation, *µ* = (ln N_2_ − ln N_1_)/(t_2_ − t_1_), where N_1_ and N_2_ are the biomass density at time t_1_ and t_2_ respectively.

### *Microcystis*-associated cultivable heterotrophic bacteria

#### Bacterial isolation

Associated bacteria were isolated from the cultures of *Microcystis* sp. on Day 16 (exponential phase of *Microcystis* sp,). Briefly, 10 mL of unialgal culture (both BG11 and BBM media) was centrifuged at 6000×g for 10 min. The supernatant was streaked on six different agar plates i.e. Muller Hilton agar, eosin methylene blue (EMB) agar, brain-heart infusion agar, Mannitol salt agar, thiosulfate-citrate-bile salts-sucrose (TCBS) agar and de Man, Rogosa and Sharpe (MRS) agar. These agar plates were incubated at 25°C for five days. On the other hand, the pellet was resuspended in 1 mL TE buffer (Tris 10 mM; EDTA 1 mM, pH 8.0). After that, 10 mL distilled water was added and sonicated three times for 3 min each time with intervals of 3 min. The solution was inoculated onto the six different agar plates as mentioned previously and incubated at 25°C for five days. Bacteria from the environment (the same date and location of the *Microcystis* sp. isolation) were also isolated using the streaking approach on same types of agars mentioned previously.

#### Identification of bacteria based on 16S rRNA amplification

Prior to 16S rRNA amplification, single colony of each bacterium was selected and cultivated in respective broth. Then, bacteria culture was subjected to genomic DNA extraction using FavorPrep Tissue Genomic DNA Extraction Mini Kit following the manufacturer’s protocols (Favorgen, Ping-Tung, Taiwan). Approximately 1500 bp fragment of the 16S gene was amplified using bacterial universal primer, 27F (forward primer) and 1492R (reverse primer). PCR amplification was performed using T-Personal Thermal Cycler (Biometra, Jena, Germany) with the same 25 µL PCR reaction as mentioned earlier except *Pfu* DNA polymerase replaced Taq polymerase. The PCR conditions were 5 min at 95°C (initial denaturation) and then 35 cycles consisting of 30 s at 95°C (denaturation), 30s at 50°C (annealing), and 1 min at 72°C (extension), and finally 7 min at 72°C for final extension. The amplified products were purified using gel/PCR DNA fragments extraction kit (Geneaid, New Taipei, Taiwan) according to the manufacturer’s protocols. Sequencing of the amplified products in bi-direction was performed by First Base Laboratories Sdn Bhd (Selangor, Malaysia).

#### Phylogenetic analysis

DNA sequence of *Microcystis* phycocyanin operon was edited and assembled separately using BioEdit version 7.1.11 software [18]. Respective references were retrieved from Genbank. The alignment of *Microcystis* phycocyanin operon was performed in this study. The DNA sequences in this study and references were aligned ClustalW implemented in the MEGA6 program package [19]. Ambiguous features (deletion, insertion or unidentified sites) were removed from the alignment. A phylogenetic tree for the alignment was constructed with maximum likelihood method based on Jukes-Cantor model, using MEGA6 program package [19]. Bootstrap values calculated from 1000 trees were utilized in the construction of phylogenetic trees. Only bootstrap values of 70 or higher were considered as significant clustering.

#### Nucleotide sequence accession numbers

The bacterial 16S rDNA, *Microcystis* 16S rDNA, and phycocyanin operon sequences have been deposited in the Genbank database under accession numbers KY794585, KY796224 and KY794583 (Table 1).

**Table 1.** Sequencing results for molecular identification of *Microcystis* sp. (isolate UPMC-A0051) using three different types of primer.

| **Cyanobacteria** | **Accession number** | **Primer** | **Reference** | **Amplified length (bp)** | **Closest relative (GenBank accession number)** | **Similarity (%)** | **Gene** |
| --- | --- | --- | --- | --- | --- | --- | --- |
| *Microcystis* sp. (UPMC-A0051) | KY796224 | CYA106F<br>CYA781R | Nubel et al. [11] | 700 bp | *Microcystis* sp. | 100 | 16S rRNA |
|  | KY794585 | PSF<br>1492R | Betournay et al. [12] | 730 bp | *Microcystis* sp. | 100 | 16S rRNA |
| | KY794583 | pcbβF<br>pcαR | Neilan et al. [13] | 685 bp | *Microcystis aeruginosa* (AF195159) | 98 | Phycocyanin |

### Statistical analysis

Differences in growth rates and normalized relative concentrations were evaluated using one-way ANOVA with Tukey *posthoc* test or two-tailed t-test. The analyses were conducted using Minitab 17 Statistical Software (Minitab Inc., State College, PA). In all cases, differences were accepted as significant when P < 0.05.

## Results and Discussion

### *Microcystis* sp. from phytoplankton blooms in Putrajaya Lake

#### Morphological characteristics

*Microcystis* sp. has been reported to be present in Putrajaya Lake previously at low density (littoral zone: 0.32%; sublittoral zone: 0.72%; limnetic zone: 0.25%) [20]. Since 2015, *Microcystis* spp. bloomed intermittently, which is different from temperate regions where blooms develop mostly during the hot weather in summer and early autumn. During the blooms, green scum was observed in the surface of Putrajaya Lake and the phytoplankton community was found to be dominated by *Microcystis* spp. The cyanobacterial species observed under a light microscope displayed irregular shaped colonies with intercellular spaces (Fig 1). The colony was composed of compact homogeneous subspherical cells with a diameter of 3.2-6.6 µm. In addition to the green scum, large number of yellow and irregular-shaped colonies with mucilaginous matrix and distal end projecting out of the colony were observed under the microscope.

**Fig 1.**
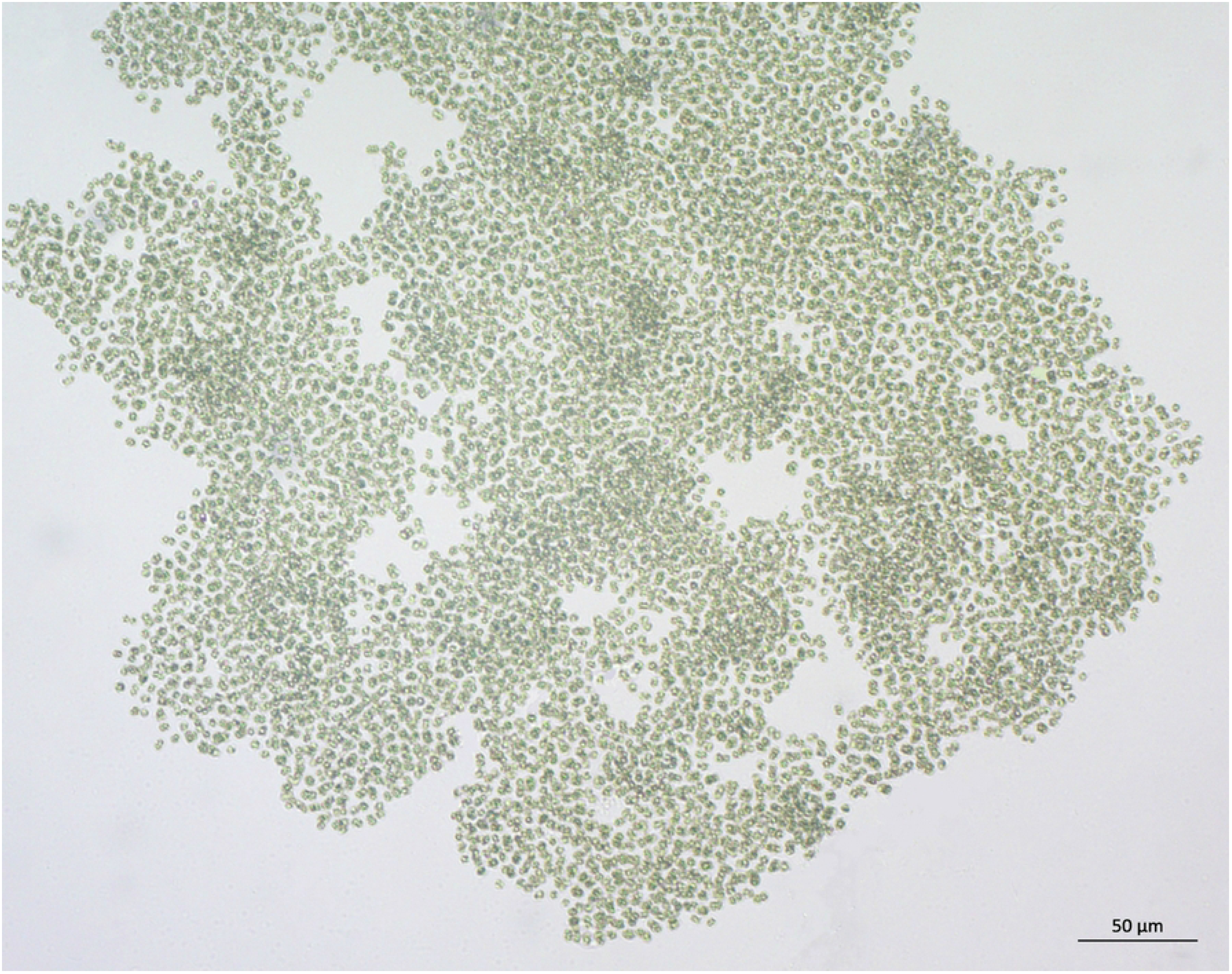
Colony of *Microcystis aeruginosa* isolated from Putrajaya Lake, Malaysia (2.9419° N, 101.6891° E) observed under a compound microscope. Scale bar = 50 µm.

#### Molecular identification

The isolated cyanobacterium from the bloom of Putrajaya Lake was identified to be *Microcystis aeruginosa* (isolate UPMC-A0051) using 16S rRNA PCR along with phycocyanin operon sequencing (Table 1). Three different sets of primer amplifying two divergent sites, 16S rRNA and phycocyanin genes (pcbβF and pcαR) used in this study produced the amplicons with amplified length ranging from 685 – 730 bp (Table 1). In comparison to some other studies that used denaturing gradient gel electrophoresis (DGGE) followed by cloning that resulted in 161 bp amplified products, longer amplified products were obtained in this study via PCR approach and direct sequencing. By obtaining a longer PCR product, more accurate and unambiguous identity could be assigned to isolated cyanobacteria. *Microcystis* spp. have been reported to be major cyanobacteria detected in different lakes of Selangor, Malaysia [20], but specific species was rarely reported.

16S rRNA PCR sequencing confirmed 100% similarity of the isolated cyanobacterium to a number of species from the genus *Microcystis*, including *M. aeruginosa, M. wesenbergii, M. novacekii, M. flos-aquae, M. ichthyoblabe, M. panniformis, M. botrys* and *M. viridis*. Phycocyanin operon sequencing is advantageous at this juncture where it showed that the isolated cyanobacterium has 98% sequence similarity with *M. aeruginosa* (Table 1). This approach demonstrated the usefulness of phycocyanin primer in assisting the confirmation of specific cyanobacterial identity. The only downside of phycocyanin primer sequencing is the limited number of phycocyanin operon sequences currently available in the Genbank. Besides, due to many 100% 16S rRNA PCR sequence similarities of the isolated cyanobacteria with references in the Genbank, phylogenetic relationship of the isolated cyanobacterium with its closest cyanobacterial reference is difficult to be illustrated. However, using phycocyanin operon sequence, the *Microcystis* in the present study could be related phylogenetically close to *M. aeruginosa* and a group of uncultured *Microcystis* sp. (Fig 2). The phycocyanin operon-based phylogeny is demonstrated to be distinct from that derived 16S rRNA gene, reflecting the fact that even organisms with 99% of similarity in the ribosomal gene may belong to different groups in terms of their pigmentation physiology. Dall’agnol et al. [21] also reported better detection of cyanobacterium microdiversity using phycocyanin operon with higher grouping capacity. *Microcystis* is a very important group of phytoplankton which is of huge medical, environmental and economic significance. Recognizing practical and efficiency phylogenetic markers of *Microcystis* is an important milestone in the development of rapid convenient kits and tools for continuous water quality analysis with high discriminatory power and capacity to yield a valid phylogenetic signal.

**Fig 2.**
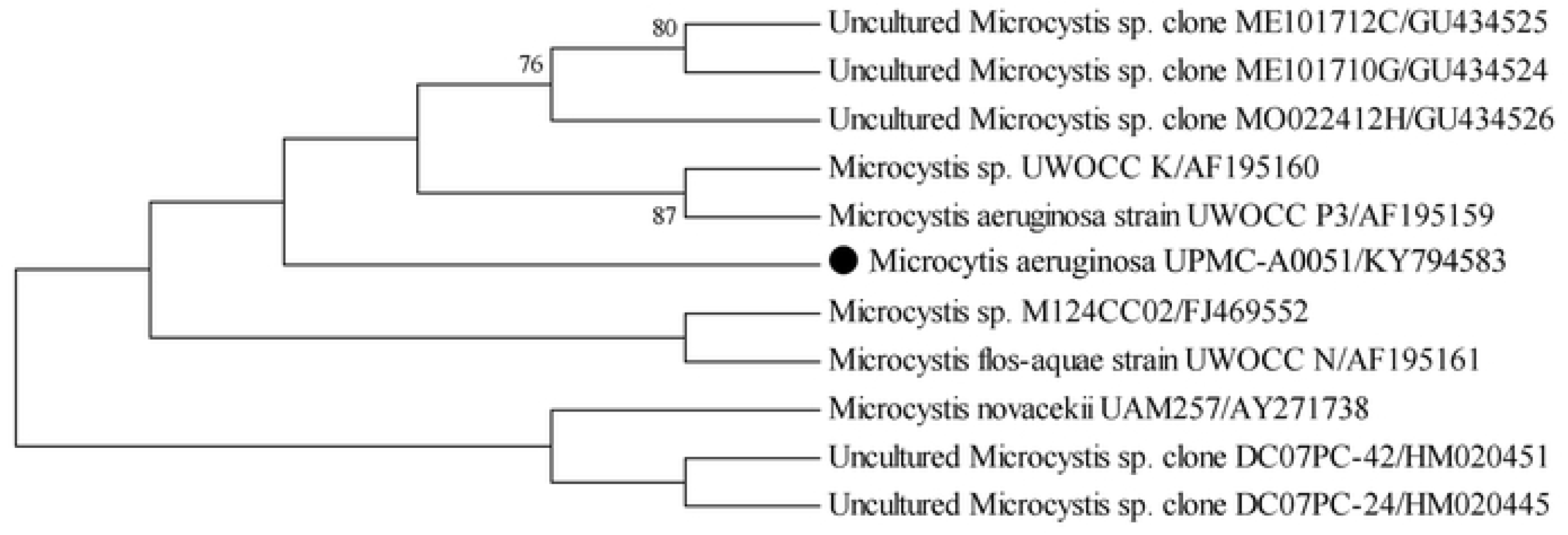
Phylogenetic tree inferred from phycocyanin operon sequences. The tree was constructed using maximum likelihood method based on Jukes-Cantor model with MEGA6 program package. The numbers above or below the internal nodes indicate bootstrap values (1000 replicates). The isolated *Microcystis aeruginosa* (UPMC-A0051) in the present study is denoted as ●.

#### Toxic producing profile

The *mcyB* A1 domain was selected as the target in the present study to analyze the toxic profile of *M. aeruginosa*. As the isolated *M. aeruginosa* (UPMC-A0051) in the present study did not contain the *mcyB* gene (Fig 3), this cyanobacterium might be a non-microcystin producing strain. Liquid chromatography tandem-mass spectrometry was further employed to confirm the presence of microcystin in the isolated strain. Chemically, the main structural variations in microcystins are observed in the L-amino acid residues X and Y, which are indicated by two-letter suffix in the name as in microcystin-LR meaning leucine (L) and arginine (R) in these positions. The molecular weight of the most common microcystin varies in the range of 909 to 1115 [22]. In this study, any compounds with a fragmentation pattern characteristic of microcystin standards was not detected within the range (Fig 4). Therefore, this implies the isolated *M. aeruginosa* (UPMC-A0051) in the present study did not belong to a microcystin-producing strain.

**Fig 3.**
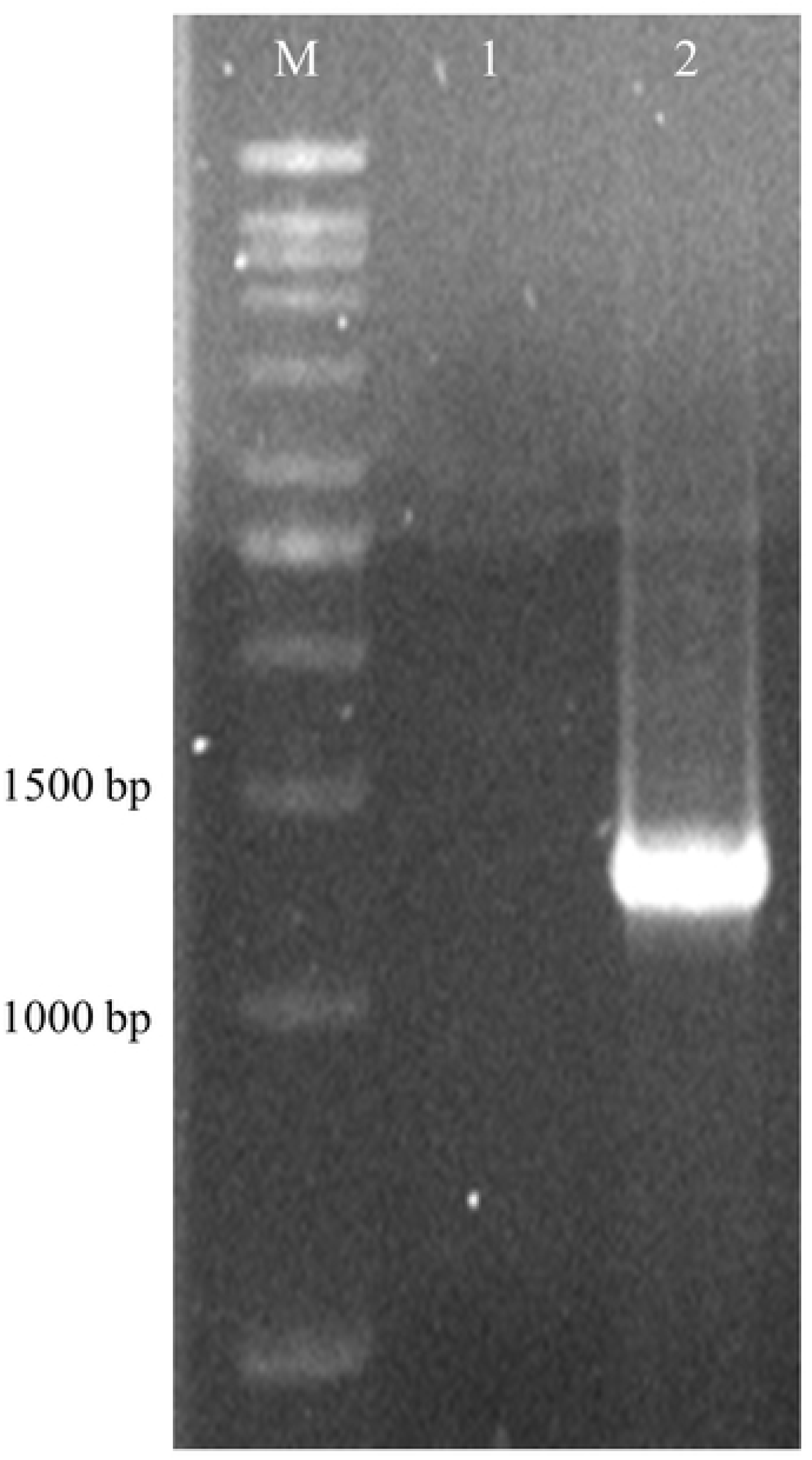
Amplification products of *mcyB* region for the isolated *Microcystis aeruginosa* (UPMC-A0051). Desired band indicating *mcyB* gene is approximately 1312 bp in size. M, Vivantis 1kb DNA ladder; 1, isolated *M. aeruginosa* (UPMC-A0051); 2, positive control (toxic *M. aeruginosa*).

**Fig 4.**
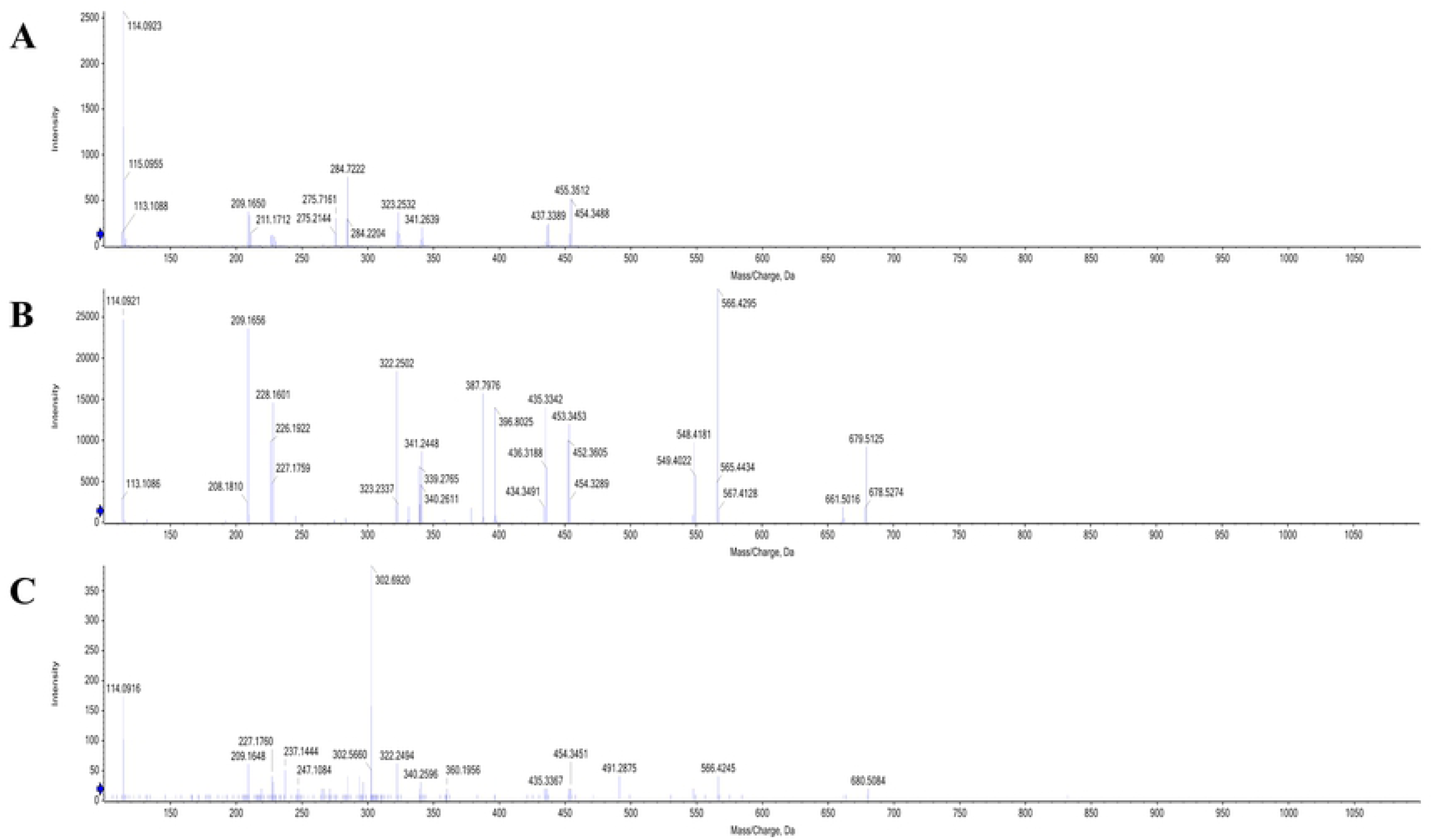
LC MS/MS profile of the isolated *Microcystis aeruginosa* (UPMC-A0051) at different retention time. (A) 5.695 min (B) 6.908 min (C) 7.410 min. The microcystins MC-RR (m/z: 520), MC-YR (m/z: 1045) and MC=LR (m/z: 995) have LC/MS retention time at 5.62 min, 6.85 min and 6.93 min, respectively.

### *Microcystis aeruginosa* (UPMC-A0051) associated cultivable heterotrophic bacteria composition in two different media and lake water

More cultivable heterotrophic bacterial species were found to grow in association with *M. aeruginosa* (UPMC-A0051) cultured in BG11 than BBM media (Table 2). Remarkably, bacteria grown in *M. aeruginosa* cultures were distinctive in different media, despite the same inoculum. Six bacterial species were discovered from BG11 and two from BBM media. Most bacteria were found to grow on Muller Hilton, brain-heart infusion and EMB media. Absence of colonies on MRS agar confirmed the absence of lactic acid bacteria such as *Lactobacillus* spp. in both the lake water and culture media. In order to differentiate between free living bacteria (planktonic) and those attached to the microalgal colony, bacteria were streaked onto the selected media agars using the supernatant and the pellet from the microalgal culture. Cogently, bacteria isolated from the supernatant of the culture represented free living bacteria; while those from the pellet were bacteria which were associated (in colony) with the cyanobacterium. Generally, all bacteria isolated from the supernatant (freely moving bacteria) were also found in the pellet (microalgal-attached bacteria) of the microalgal culture. Two bacteria were isolated from BBM media and identified as *Sphingomonas parapaucimobilis* (M1) and *Hydrogenophaga intermedia* (M2) (Table 3A). Both of them were found present in the medium as free-moving and attached bacteria. Meanwhile, six bacteria were isolated from BG11 medium in association with *Microcystis aeruginosa*. Of the six bacteria isolated from BG11 culture, three bacteria were found within both the supernatant and the pellet. Those three were identified as *Microbacterium oxydans* (G1), *Blastomonas natatoria* (G2) and *Ochrobactrum* sp. (G3). Three other bacteria isolated from the BG11 culture were found only from the pellet, denoting their attachment to *M. aeruginosa.* These were *Microbacterium hydrocarbonoxydans* (G4), *Microbacterium trichothecenolyticum* (G5) and *Microbacterium hominis* (G6) (Table 3A). This is in line with Shen et al. [4] who reported that some *Microcystis* species harbour heterotrophic bacteria within their large mucilaginous colonies that might be responsible for the aggregation of cyanobacterial cells.

**Table 2.** Summary of bacteria from both pellets and supernatant from two different media on agar plate. Bacteria from Bold’s Basal Medium (BBM) and BG11 culture media were cultivated on agar media, coded as M1-M6 and G1-G6 for BBM and BG11, respectively.

| Agar | BBM |  | BG11 |  |
| --- | --- | --- | --- | --- |
|  | Pellet | Supernatant | Pellet | Supernatant |
| Muller Hilton | M1 | M1 | G1 | G1 |
|  |  |  | G2 | G2 |
|  | M2 | M2 | G3 | G3 |
|  |  |  | **G4** |  |
|  |  |  | **G5** |  |
| Brain-heart infusion | M1 | M1 | G1 | G1 |
|  |  |  | G3 | G3 |
|  |  |  | **G6** |  |
| EMB | M2 | M2 | G1 | G1 |
|  |  |  | G2 | G2 |
|  |  |  | G3 | G3 |
|  |  |  | **G5** |  |
| TCBS | — | — | G1 | G1 |
| Mannitol Salt | — | — | G1 | G1 |
| MRS | — | — | — | — |

Bold denotes isolated bacteria that only found in pellet but not supernatant.
EMB, Eosin methylene blue; TCBS, Thiosulfate-citrate-bile salts-sucrose; MRS, de Man, Rogosa and Sharpe.

**Table 3.** 16S rRNA PCR identification of *Microcystis*-associated bacteria isolated from defined media and environment. (A) Defined media (BG11 and BBM); and (B) Putrajaya Lake. G1-G6, M1-M2 and E1-E16 are bacteria isolated from BG11, BBM media and the Putrajaya Lake, respectively.

| Media | Isolated bacteria | Closest relative (GenBank accession number) | Similarity (%) |
| --- | --- | --- | --- |
| BG11 | G1 | *Microbacterium oxydans* (JN644529) | 100 |
|  | G2 | *Blastomonas natatoria* (HF930749) | 100 |
|  | G3 | *Ochrobactrum* sp. (KJ957921) | 100 |
|  | G4 | *Microbacterium hydrocarbonoxydans* (KR052878) | 99.8 |
|  | G5 | *Microbacterium trichothecenolyticum* (KJ631291) | 100 |
|  | G6 | *Microbacterium hominis* (HM032799) | 99.8 |
| BBM | M1 | *Sphingomonas parapaucimobilis* (AB680768) | 100 |
|  | M2 | *Hydrogenophaga intermedia* (KT719641) | 99.9 |

**B.**
| Isolated Bacteria | Closest relative (GenBank accession number) | Similarity (%) |
| --- | --- | --- |
| E1 | *Aeromonas* sp. | 100 |
| E2 | *Aeromonas* sp. | 99.9 |
| E3 | *Exiguobacterium* sp. (KC688873) | 100 |
| E4 | *Exiguobacterium* sp. | 100 |
| E5 | *Enterobacter* sp | 100 |
| E6 | *Vogesella perlucida* (HE614874) | 99.9 |
| E7 | *Pseudomonas* sp. | 100 |
| E8 | *Stenotrophomonas maltophilia* (KT986130) | 100 |
| E9 | *Elizabethkingia miricola* (EU375848) | 99.8 |
| E10 | *Delftia litopenaei* (NR108843) | 100 |
| E11 | *Bacillus* sp. | 100 |
| E12 | *Acinetobacter* sp. | 100 |
| E13 | *Pantoea dispersa* (KU597506) | 100 |
| E14 | *Pseudomonas alcaligenes* (KT336735) | 100 |
| E15 | *Acidovorax* sp. | 99.9 |
| E16 | *Pseudomonas* sp. | 100 |

Bacteria were postulated to come from within *Microcystis* mucilage in the form of cells as well as spores. Deductively, there should be some kind of similarity in the bacteria composition within cultures of cyanobacterium in BG11 and BBM given similar inoculum. However, such was not true in this study as two distinct compositions of cultivable heterotrophic bacteria associated with *M. aeruginosa* in BG11 and BBM media were observed. Besides, compared to the total of 16 types of bacteria isolated from Putrajaya Lake (Table 3B), none of the eight bacteria from *M. aeruginosa* grown under the controlled environment (two from BBM and six from BG11) were similar. These eight bacteria isolated from *M. aeruginosa* cultured in BBM and BG11 could be classified into three different classes, *Actinobacteria, α.-Proteobacteria* and *β-Proteobacteria*. These bacterial classes were found in cyanobacteria culture and lake cyanobacterial blooms using culture dependent and culture independent approaches [23,24]. In addition, bacteria from *Actinobacteria* and *Protobacteria* classes have been reported to interact with *M. aeruginosa* in BG11 medium at 25°C [4]. However, different genera of bacteria were obtained in the present study although the same BG11 medium was utilized to culture the *M. aeruginosa*, compared to the previous study. Amongst the *Microcysti*s-associated bacteria found here, *Microbacterium* spp. (which represented the majority of the bacteria found in association with *M. aeruginosa* cultured in BG11 media in this study), *Ochrobacterium* sp. and *Sphingomonas* sp. have been reported to be capable of utilizing and degrading microcystin [25]. *Hydrogenophaga* has been reported to be an algicidal bacterium against harmful algae bloom [9]. These *Microcysti*s-associated bacteria except *Ochrobacterium* and *Hydrogenophaga* have been observed in lake cyanobacterial blooms using culture-dependent method [23]. Shi et al. [26] showed that *Microcystis*-associated bacteria were naturally selected by means of toxin tolerance and/or ability to utilize exudates from *Microcystis*. This study considers the latter to be more acceptable, considering the isolated *M. aeruginosa* (UPMC-A0051) in this study has been demonstrated to be incapable of producing toxic microcystin, thus subtracting the conjecture that symbiotic bacteria within the cultures did not arise due to elimination or tolerant factor of the bacteria to toxins. Therefore, composition of cultivable heterotrophic bacteria interacting with *Microcystis* were found to be more motivated by food/nutrient factor and the microalgae-bacteria interaction was not microalgae-species or bacteria-species specific. It is postulated that it is more possible that it is the unique biochemical signature produced by *Microcystis* spp. that acts as the contributing factors to the types of bacteria associating with it. Fuhrman et al. [27] reported the marine bacteria community was mainly affected by water temperatures. Perchance, different sampling regions such as temperate and tropical areas may yield different bacterial composition. A tropical *M. aeruginosa* isolate may also exhibit varied properties and adaptation traits in its symbiotic interactions and survival.

Chemically, BG11 and BBM media shared similar elements, albeit in varying composition. Distinctively, sodium nitrate is approximately six times higher in BG11 as compared to the BBM medium. It is very interesting to note that sodium nitrate is a common preservative used in meat products and higher concentration of sodium nitrate in BG11 also gave rise to more bacterial species in this media compared to BBM. It could be possible that sodium nitrate boosted microalgae growth rate that in turn led to increased secretion of microalgal byproducts that might have promoted the bacterial growth. On the other hand, BBM medium included a buffering system consisting of K_2_HPO_4_ and KH_2_PO_4_, favouring to an acidic solution, as well as nearly 7.5 times higher in potassium as compared to BG11. Earlier studies have indicated toxicity of potassium towards *Microcystis* spp. [28,29] but its toxicity towards heterotrophic bacteria is less known. Again, it is notable in this study (in comparison with earlier study [4]) that even among similar culture media (i.e. BG11), dissimilar genera of bacteria would be found associating with similar microalgae (i.e. *M. aeruginosa*). This seems to be in line with Sapp et al. [30], that inorganic nutrient level has no impact in shaping a bacterial community. On the other hand, Brunberg [31] demonstrated that exopolysaccharides (mucilage) derived from *Microcystis* might create an appropriate microenvironment for the bacteria to form biofilm that may play a role in modifying the relationship between cyanobacteria and bacteria [32]. The present study demonstrated that a culture environment (defined as culture media) directly affects possible cultivable heterotrophic bacteria composition growing in association with *M. aeruginosa*. It might be possible that different media would affect the production of *Microcystis* endo- and exopolysaccharides, which indirectly influenced the bacterial composition. Overall, the isolated bacteria in the present study all appeared to be capable to utilizing products and byproducts of *Microcystis*, highlighting possible food chain link with *M. aeruginosa*. Further studies are needed to elucidate the factors that influence the bacterial community associated with microalgae.

### Influence of varying bacterial composition in different media on the morphological and physiological characteristic of *M. aeruginosa*

#### Growth characteristics of *M. aeruginosa* are affected by the bacterial composition within the growth media

Generally, *M. aeruginosa* (UPMC-A0051) showed significantly higher (p < 0.05) biomass growth when cultured in BG11 compared to BBM media (Fig 5A). As compared to their axenic counterpart, *M. aeruginosa* grown in association with the bacteria showed significantly higher biomass production (p < 0.05). The microalgae showed continual growth-decline pattern (in terms of biomass) in every 4-5 days on average. The biomass of microalgae only showed sign of plateau on Day 26 till the end of the study on Day 30 recording 0.32-0.40 g/ L and 0.39-0.46 g/ L for those grown in BBM and BG11, respectively. The highest biomass for BG11 and BBM media was observed on Day 24, with values of 0.49 g/ L and 0.44 g/ L, respectively. It is not uncommon to observe different growth rates within a batch culture of microalgae in an extended culture period. Previously, Krzeminska et al. [33] showed two distinct growth phases in five green microalgae cultured for 10 days where the first growth phase involved Day 0 to Day 3. However, it is most peculiar to observe repeating sets of biomass rise and decline in an almost fixed amount of time. As the same observation was noted similarly in both media, this pattern of continuous biomass oscillations was most probably attributed to the bacteria living symbiotically with the microalgae, where nutrient degradation and recycling were most probably involved. Comparatively, it was found that *M. aeruginosa* cultivated in BG11 medium showed significantly higher biomass per volume compared to that cultured in BBM medium throughout the whole culture period (p < 0.05). This pattern also coincided with the richer bacterial composition found in BG11 culture than BBM, as discussed previously. It is hypothesized that richer bacterial species composition could aid in nutrient degradation and recycling more efficiently, hence led to a higher biomass per volume of *M. aeruginosa*.

**Fig 5.**
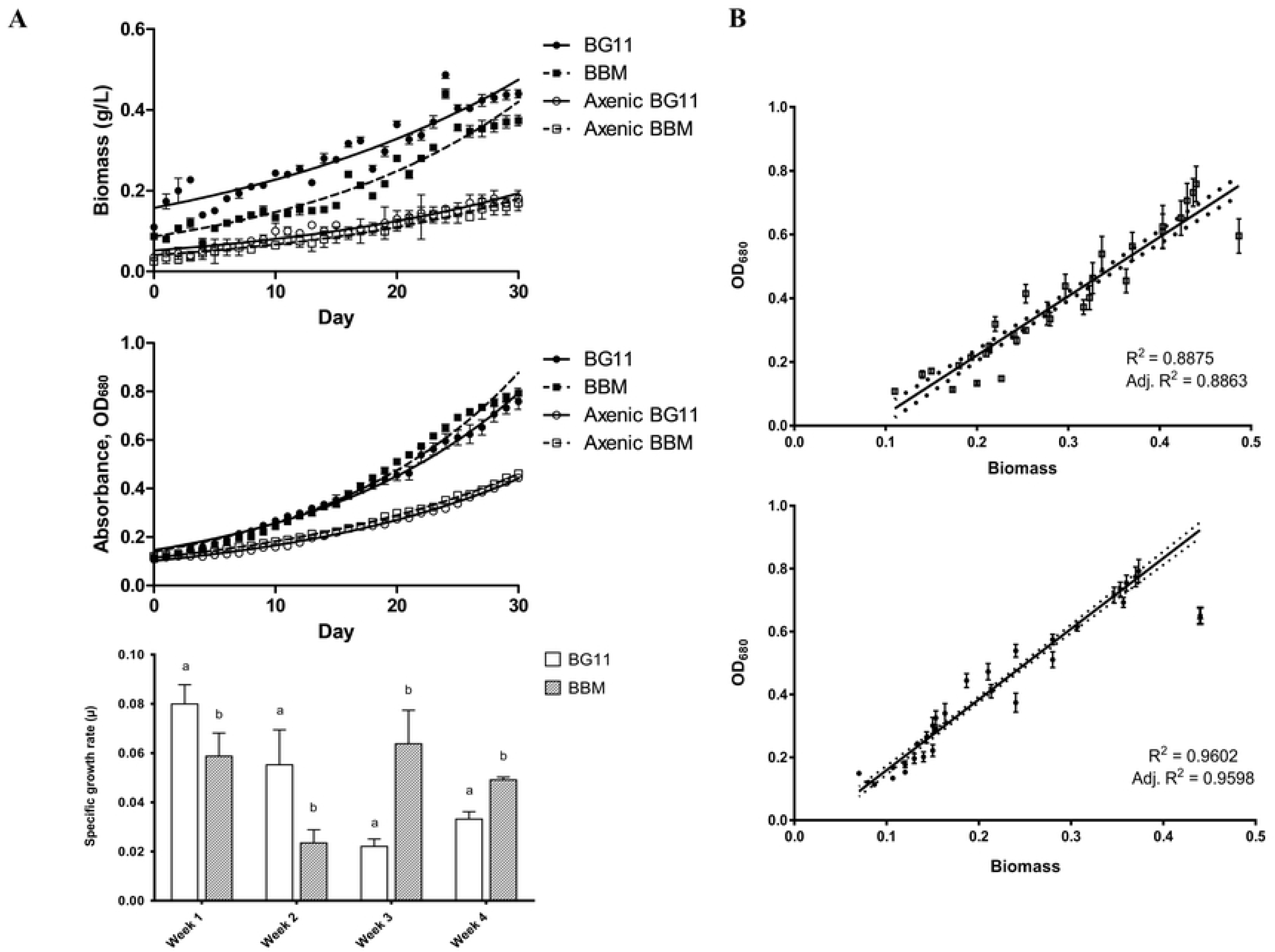
Growth properties of *Microcystis aeruginosa* (UPMC-A0051). (A) Growth characteristics of *Microcystis aeruginosa* (UPMC-A0051) grown in BG11, axenic BG11, BBM and axenic BBM media, measured as dry biomass (g/L), optical density at 680 nm and specific growth rate for 30 days. Dotted line denotes best-fit curve plotted using least-square regression for the estimation of growth rate. Values represent means (n=3) and error bars represent standard deviations. (B) Correlation between algal biomass (g/L) and optical density (OD_680_) of *Microcystis aeruginosa* (UPMC-A0051) in BG11 (□) and BBM (•) media. Values are presented as means (n=3) with R^2^ represents correlation coefficient and error bars represent standard deviation.

On the other hand, indirect measurement of cell growth using optical density quantification at 680 nm showed that *M. aeruginosa* cultured in BBM medium demonstrated slightly higher growth than BG11 although microalgae grown in both media showed continuous increase over the period of 30 days (Fig 5A). Similarly, *M. aeruginosa* grown in association with the bacteria showed significantly faster colour development as compared to their axenic counterpart (p < 0.05). Statistically, the estimated growth rate (in best-fit values) for *M. aeruginosa* cultured in BG11 and BBM for 30 days was 0.025 and 0.027, respectively. As the values of OD_680_ reflect chlorophyll *a* concentration and cell density of the culture, cyanobacteria grown in BBM medium was found to develop the photosynthetic pigment slightly faster than that grown in BG11. Rather apparent was the colour displayed by the cultures in BG11 and BBM broth media which was blue green and bright olive green, respectively. Besides, Shukla and Rai [29] reported that higher potassium concentration would lead to increased carotenoid content in *M. aeruginosa*, striking visual differences in overall medium colour probably due to variations in carotenoid profiles. This may presumably contributed by the cyanobacteria’s interaction with the different bacteria grown in each media. Interestingly, *M. aeruginosa* grown in media harbouring higher number of bacterial species, i.e. BG11, also showed lower growth correlation with OD_680_, as compared with those grown in BBM (that have lower number of bacterial species) (Fig 5B). The coefficient of determination of algal biomass against optical density (OD_680_) of *Microcystis* spp. cultivated under axenic condition has been reported in many studies to be 0.95 and higher [34]. This study also suggested that microalgae growth assessment by means of optical density might be affected by the axenicity of the microalgae culture.

*Microcystis aeruginosa* (UPMC-A0051) exhibited higher specific growth rate (SGR) in BG11 for the initial two weeks while those grown in BBM showed higher growth rates from Week 3 (Day 14-21) till the end of the study (Day 21 to 30) (Fig 5A). Most possibly, the SGR pattern quantified in this study was highly influenced by the interacting bacteria within the same medium. Competition may be the most possible cause of decreasing SGR in BG11-cultured *Microcystis aeruginosa*, which may also be true to BBM-cultured *Microcystis*, as the BG11 was found to harbour higher number of bacterial species earlier as compared to BBM. In addition, bacteria growing in BBM were found to be free-living as compared to BG11 where half of the bacteria species were found to be attached to the *Microcystis* biomass. Eventually, the possibility of bacterial behaviour in affecting specific growth rate of microalgae was indicated in this study.

### Morphological and colony characteristics of *M. aeruginosa* varied among different bacterial composition

*Microcystis aeruginosa* (UPMC-A0051) grown in BG11 medium was found to multiply and formed colonies faster than that cultivated in BBM medium (Fig 6). The microscopic observations concurred well with the findings on the biomass and pigment development of *Microcystis aeruginosa* as discussed before.

**Fig. 6.**
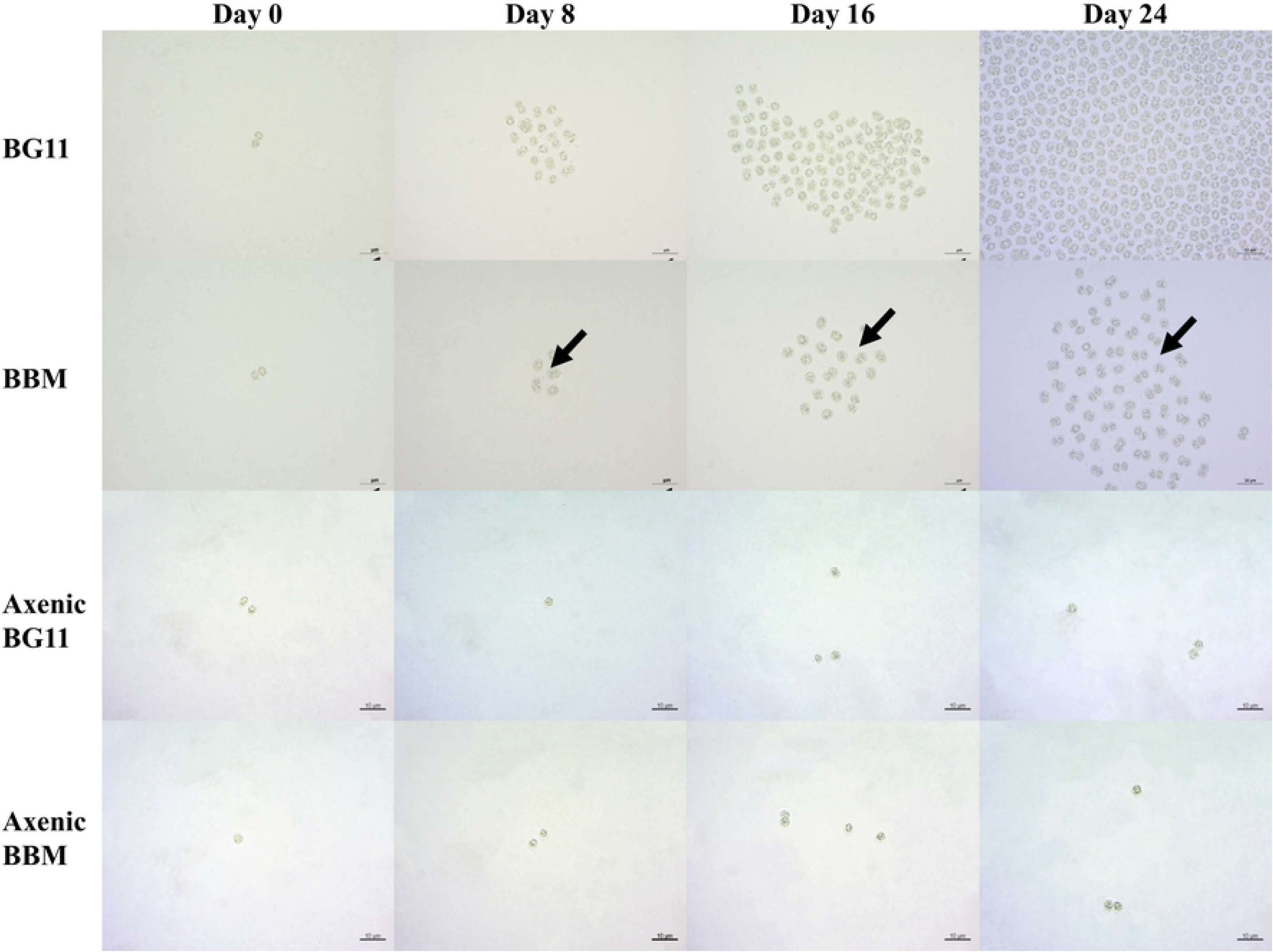
Morphological features of *Microcystis aeruginosa* (UPMC-A0051) in BG11, axenic BG11, BBM and axenic BBM media on different days. Arrows show the space between solitary cells within the clathrate of *M. aeruginosa*, which is more apparent in colonies grown in BBM medium (Scale bar = 10 µm).

Initially, *M. aeruginosa* inoculated into both BG11 and BBM media showed similar morphological features (Day 0). The solitary cell of the cyanobacteria in this study was observed to consist of, in majority, a pair of spherical (some slightly elongated) cells, about 2.0 μm in diameter each. Subsequent observations, however, found that the cyanobacteria in different media slowly showed distinct morphological features especially on their colonization and mucilagous formation. Most obviously, there was a difference between *M. aeruginosa* grown in association with the bacterial composition with those grown in axenic culture. Particularly, *M. aeruginosa* grown in axenic culture were observed to hardly form colonies as compared to those in association with bacteria, where it was also observed that the cyanobacteria exhibited different colony characteristics under different media.

It is interesting to note that the space between solitary cells within the clathrate of *M. aeruginosa* grown in BBM media was found to be more intense that those grown in BG11, and this space increased with the growing period (Fig 6). *Microscystis aeruginosa* in BG11 culture displayed a firmly bound colony, indicating smaller spaces in between cells in a colony, similar to the *M. aeruginosa* collected from the environment (Fig 1). This space was found to be consistent, about 1-2 µm apart, along the observation period of 24 days. On the other hand, *M. aeruginosa* colonies in BBM culture was found to be loosely bound, indicating bigger spaces in between cells (3-8 µm apart) and higher volume of mucilage. This coincided well with the bacterial community study aforementioned that polysaccharide production (in the form of mucilages) of the cyanobacteria was highly influenced by different culture media that would in turn influence the structure of bacterial community growing in interaction with the cyanobacteria, and vice-versa.

Earlier, the morphological characteristics of *M. aeruginosa* grown axenically in BG-11 medium has been illustrated where the cyanobacteria typically developed from solitary cells to compact colonies followed by daughter colony segregation; and finally clathrate colony after 8–10 days of growth [34]. Observation of *M. aeruginosa* grown in this study was found to be slightly different. Besides, *M. aeruginosa* colonies were found to grow and develop faster in BG11 medium, which has been found to harbour higher number of bacteria species than those in BBM. *Microcystis* cultured in BBM also exhibited lesser number of cells within a clathrate compared to that cultured in BG11. Shen et al. [4] reported that the presence of heterotrophic bacteria induced the aggregation of *M. aeruginosa* cells into colonies. The present study also noticed that heterotrophic bacteria were important components in the colony formation of *M. aeruginosa*. A higher number of bacterial species associated with *M. aeruginosa* were noticed in BG11 compared to BBM media. It is postulated that the presence of higher number bacterial-associated *M. aeruginosa* in terms of species (BG11 media) were likely to “squeeze” the *M. aeruginosa* cells into compact colonies without any spaces. These results probably support the slight difference of colony pattern of *M. aeruginosa* in these two culture media. Therefore, it could be concluded as *Microcystis*-associated bacterial composition aid in the aggregation of *Microcystis* cells into particular pattern of colonies to enhance its adaptation in different environment.

## Conclusion

The isolated cyanobacterium from the phytoplankton bloom of Putrajaya Lake was identified to be *Microcystis aeruginosa* (isolate UPMC-A0051). None of the bacteria isolated directly from the lake were similar to the bacteria associated with the *M. aeruginosa* grown in BG11 and BBM media. Significantly higher number of bacterial species was found in BG11 than BBM media. *Microcystis aeruginosa* grown in BG11 medium also showed significantly higher accumulation of biomass compared to that in BBM medium (p < 0.05). The morphology of *M. aeruginosa* in both media was also found to be different where cells in colonies cultured in BG11 medium were more compact compared to those in BBM medium. As compared to their axenic counterpart, *M. aeruginosa* grown in association with the bacteria showed significantly higher biomass production (p < 0.05). This study illustrated that *Microcystis*-associated bacterial composition could enhance the cyanobacterial growth and aid in the aggregation of *Microcystis* cells into particular pattern of colonies. Further studies should be performed using the isolated heterotrophic bacteria to understand the role of these bacteria in shaping the morphological and physiological of *Microcystis aeruginosa* UPMC-A0051.

## Acknowledgements

This study was supported by the Ministry of Education Malaysia under the Fundamental Research Grant Scheme (FRGS) Project No. FRGS/2/2014/STWN01/UPM/01/1 and SATREPS-COSMOS matching fund for the Malaysia-Japan SATREPS-COSMOS (JICA-JST) collaborative program. Technical assistance from staff and students of the Laboratory of Marine Biotechnology, Institute of Bioscience is duly acknowledged.

